# Genome sequence and description of *Blautia brookingsii* str SG772 nov., a new species of anaerobic bacterium isolated from healthy human gut

**DOI:** 10.1101/327007

**Authors:** Sudeep Ghimire, Roshan Kumar, Eric Nelson, Jane Christopher-Hennings, Joy Scaria

**Affiliations:** Department of Veterinary and Biomedical Sciences, South Dakota State University, Brookings, SD, USA.; South Dakota Center for Biologics Research and Commercialization, SD, USA

**Keywords:** *Blautia*, *brookingsii SG772*, Human gut, Genome, Anaerobe

## Abstract

Strain SG-772 is a Gram positive, strictly anoxic bacterium isolated from the feces of a healthy human fecal donor. Based on 16S rRNA gene sequence, the strain showed maximum similarity (94.39%) with *Blautia stercoris* GAM6-1 in EZ-Taxon server and thus assigned the genus *Blautia*. The scanning electron micrograph of the bacterium revealed the characteristic coccobacillus shape as well as the complete absence of flagellum, suggesting its non-motile phenotype. This strain was found to utilize 27 substrates based on Biolog AN plates assay, with maximum preference for D-mannitol. Additionally, the strain was found to be resistant to tetracycline and streptomycin. Genome sequencing and analysis revealed an overall genome size of 3.49 Mbp and GC content of 43.97%. Based on RAST annotation server, the closest neighbor was *Blautia hansenii* DSM20583. Average Nucleotide Identity (ANI) of these strains were 81.69%, suggesting a high level of genomic variation. The comparative genome analysis of strain SG772 with *B. hansenii* DSM20583 revealed a total of 411 orthologous genes coding for basic metabolic functions. Furthermore, the genomes were functionally distinct based on COG categories. Thus, based on all these differences, we propose a novel species of genus *Blautia* named as *Blautia brookingsii* SG772.

## Introduction

The members of family *Lachnospiraceae* are commensal bacteria inhabiting the human gastrointestinal tract (1). These group of bacteria degrade complex polysaccharides to produce short chain fatty acids. Acetate, butyrate and propionate produced as the result of digestion of polysaccharides can be used as an energy source for the host (2). *Lachnospiraceae* in human gut microbiota plays important role in maintaining health and disease balance (3–7). Recently, a new genus *Blautia* has been formed under *Lachnospiraceae* family by reclassifying few members of genus *Clostridium* and *Ruminococcus* (8). *Blautia* has been reported to provide colonization resistance against *Enterococcus faecium* (9) and *Clostridium difficile* (5).

The members of genus *Blautia* are gram-positive, non-motile, coccoid shaped; obligate anaerobes which can produce a wide variety of compounds as the by-product of fermentation. Most of the members of this genus were either isolated from the human or mammalian fecal samples (8, 10). During the culturomics of human feces, a novel *Blautia*-like strain was isolated from Brain Heart Infusion (BHI) based medium and named as strain SG772. Based on 16S rRNA gene sequence phylogeny, it was taxonomically clustered with the genus *Blautia*. Additionally, using taxo-genomics approach, the bacterium SG772 showed distinct morphological, physiological and genomic attribute as compared to its closest neighbor. Based on all these differences, we propose a novel species of genus *Blautia* named as *Blautia brookingsii* SG772.

## Organism information

### Growth conditions and biochemical properties

During culturomics of healthy human fecal sample, strain SG772 was isolated from a healthy human fecal sample using Brain Heart Infusion (BHI) agar medium anaerobically at 37°C. Morphologically, colonies were round, whitish, convex and smooth after 48 hours incubation on BHI agar. Further, cellular morphology of strain SG772 was examined using a scanning electron microscope. Microscopic study revealed the complete absence of flagellum, indicating that strain SG772 is non-motile. The size of the bacterium varied between 0.5–0.8 × 1.8–2.5 μm (Figure 1). The Gram-stain test was performed using a Gram staining kit (HiMedia) and the strain was found to be Gram-positive. Optimum temperature and pH for growth of the bacteria was found to be 37°C and 6.8 respectively.

**Figure 1:**
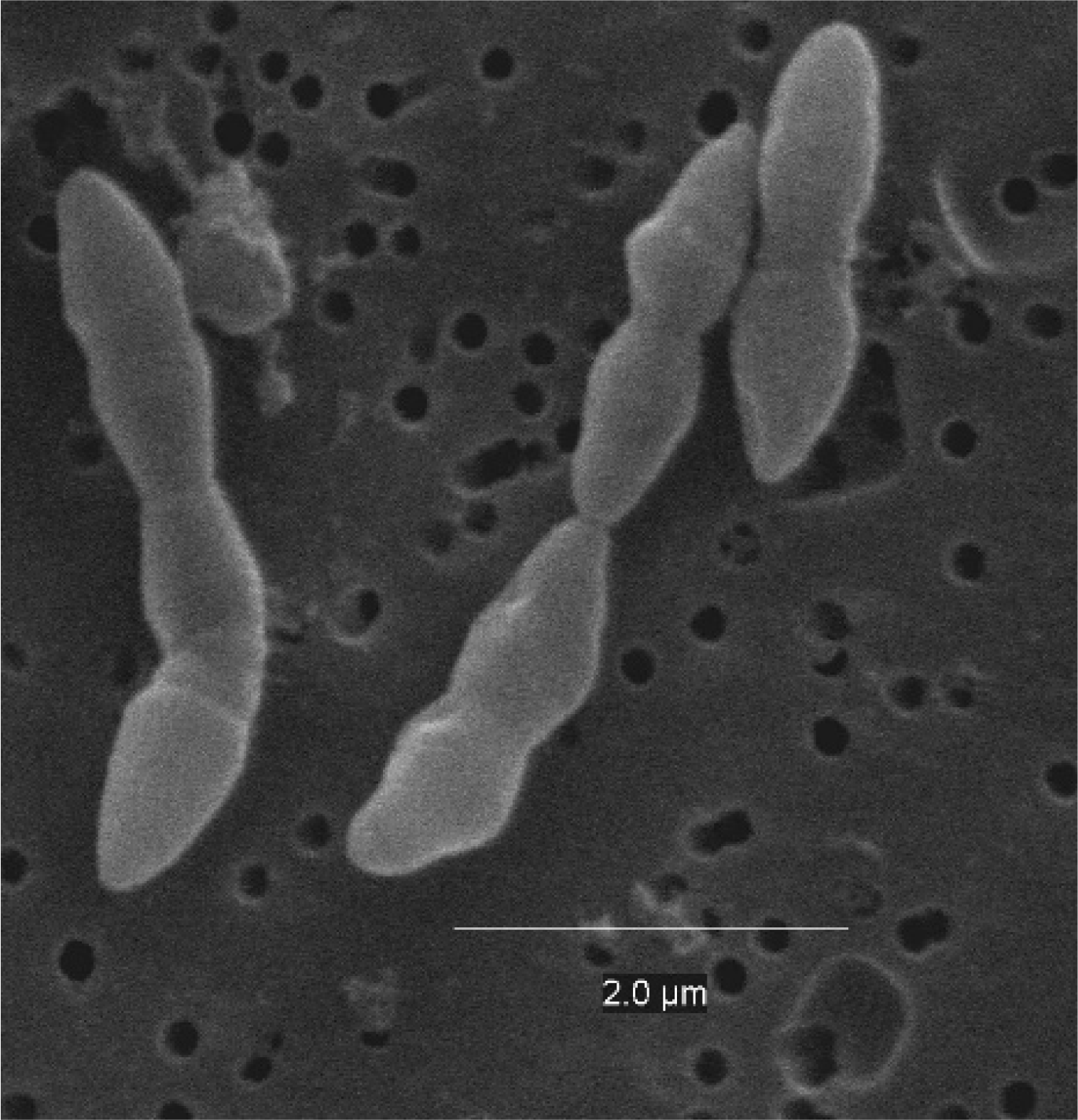
Scanning electron micrograph of strain SG772 grown on BHI agar at 37°C for 24 hours. Bar, 2.0 μm

As the strain SG772 exhibited no significant scores on Matrix-assisted desorption-ionization time-of-flight (MALDI-ToF) (Bruker Daltonics, Germany) (11), the genes encoding for 16S rRNA gene was amplified, sequenced using Applied Biosystems 3500xL, Genetic analyser (Applied Biosystems, MA, USA) at Department of Veterinary and Biomedical Sciences, South Dakota State University. The sequence was assembled using Sequencing Analysis version 5.1.1 Genious 10.2.3 (NJ, US). A continuous stretch of 1440 bp of the 16S rRNA gene of strain SG772 was obtained and this sequence was subjected to similarity search against EzTaxon-e-server (12). Analysis of 16S rRNA gene revealed highest sequence similarity (94.39%) to *Blautia stercoris* GAM6-1. Thereafter, the phylogenetic tree was constructed based on 16S rRNA gene sequence, together with top 30 taxonomically characterized strains. The evolutionary distance matrix was calculated using the distance model of Jukes & Cantor (1969) and an evolutionary tree was reconstructed using the neighbor joining method. The sequence of *Atobobium minutum* NCFB2751 was used as an outgroup (Figure 2). The resultant tree topology was evaluated by bootstrap analysis based on 1000 replicates (Felesenstein J, 1985) in Mega version 5.2.2. Strain SG772 falls in the clade containing members belonging to the genus *Blautia*. (Figure. 2)

**Figure 2:**
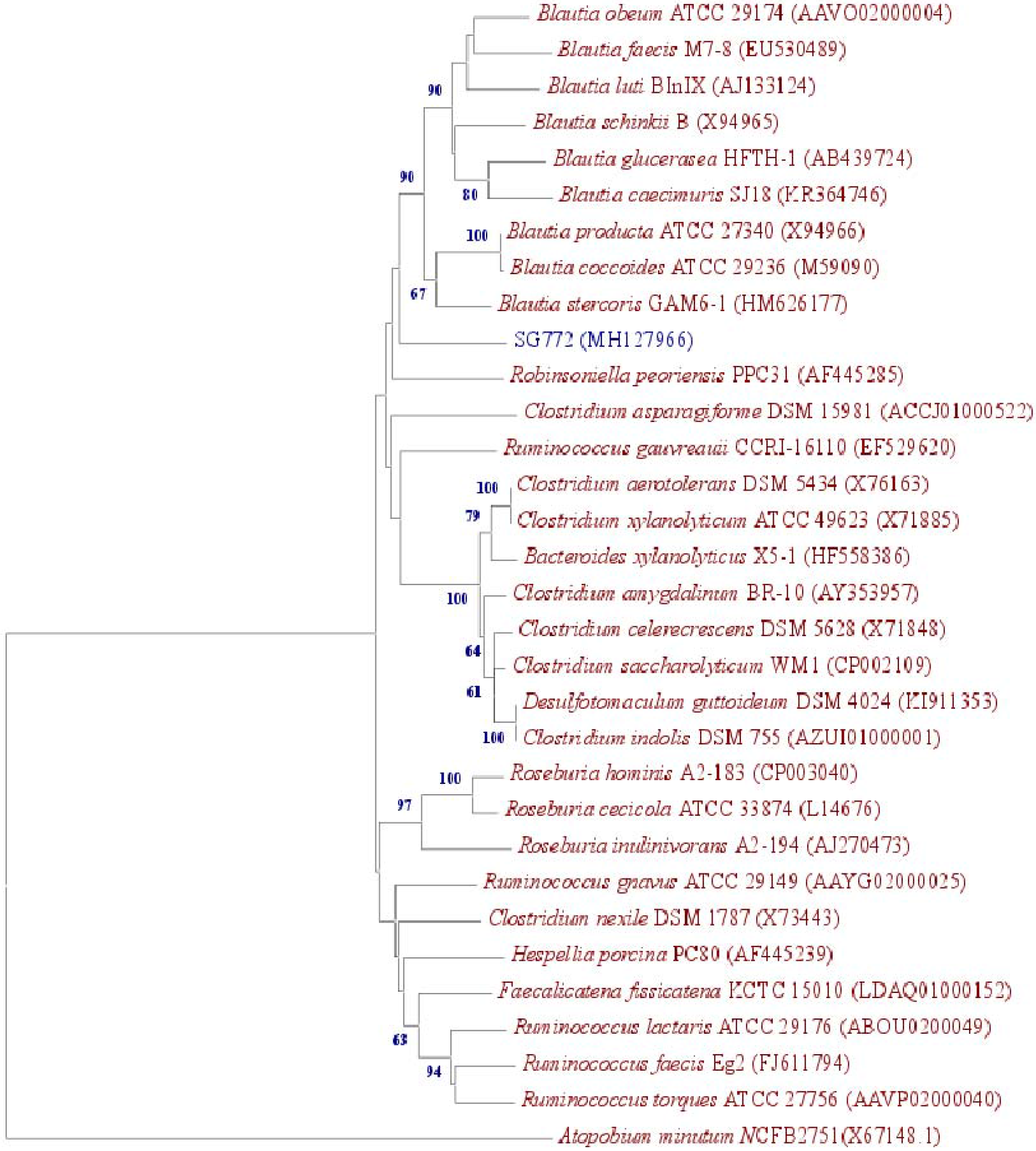
16s rRNA sequence based phylogenetic tree constructed using neighbor-joining method. The 16S rRNA gene sequence was compared against the EZ-taxon, which showed the highest similarity (94.39%) with *Blautia stercoris* GAM-61. The sequence of *Atobobium minutum* NCFB2751 was used as an outgroup.Numbers at nodes indicate bootstrap value expressed as percentages of 1000 replications. Bar, 0.02 accumulated changes per nucleotide. GenBank accession numbers are shown in parentheses.

The growth pattern for the strain SG772 was determined using growth curve assay (Figure 3A). The biphasic growth pattern indicate its ability to use simple sugars in media initially with utilization of complex carbohydrates when simple sugars are depleted. Additionally, in order to elucidate the pattern of substrate oxidation, Biolog assay was performed using Biolog AN MicroPlate^TM^ (Biolog Catalog # 70007) anaerobically. For this, 100 μl of overnight grown culture was plated onto 150 mm BHI agar plate and incubated for 48 hours anaerobically at 37° C. Grown colonies were picked by a sterile cotton swab and inoculated to AN inoculating fluid (Biolog Catalog # 72007) until OD_650_ reached 0.3. From this suspension, 100μl was pipetted into each well of 96 well biolog AN microplate in triplicate and incubated at 37°C anaerobically. OD_650_ readings were taken at 0 hour and 24 hours post inoculation and results were analyzed and compared against *Blautia stercoris* GAM6-1 (Table 1). The strain was found to utilize 27 different substrates with maximum preference for D-mannitol (Figure 3B).

**Figure 3:**
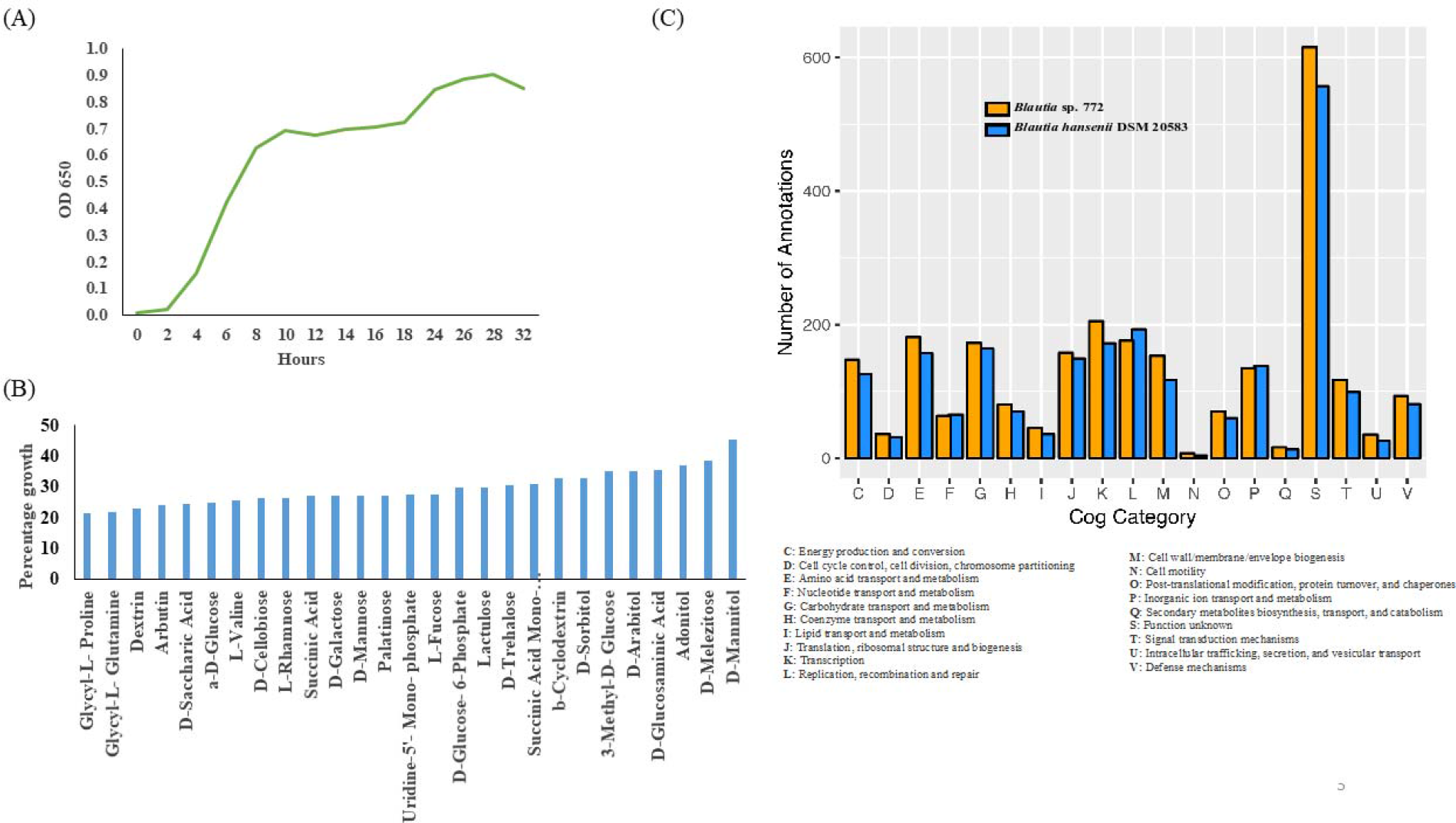
Phenotypic and genotypic profiling of *Blautia sp.* SG772-. (A) Growth curve of the strain *Blautia sp.* SG772, (B) Bar plot depicting different patterns of substrate utilization, (C) Bar graph depicting comparative abundance of COG gene families between the two genomes.

**Table 1:** Physiological features of strain SG-772 compared to phylogenetic neighbor *Blautia stercoris* GAM6-1. (*Data adopted from Park et al., 2012)

| Characteristics | SG772 | *Blautia stercoris* GAM6-1* |
| --- | --- | --- |
| Habitat | Human gut | Human gut |
| Classification | Domain: Bacteria | Domain: Bacteria |
|  | Phylum: Firmicutes | Phylum: Firmicutes |
|  | Class: Clostridia | Class: Clostridia |
|  | Order: Clostridiales | Order: Clostridiales |
|  | Family: Lachnospiraceae | Family: Lachnospiraceae |
|  | Genus: *Blautia* | Genus: *Blautia* |
|  | Species: *brookingsii* | Species: *stercoris* |
| Optimum temperature | 37°C | 37°C |
| Shape | Coccobacilli | Cocci |
| Optimum pH | 6.8 | 6.2 |
| α-Galactosidase activity | - | + |
| β-Galactosidase activity | - | + |
| α-Glucosidase | - | + |
| β –Glucosidase | - | + |
| N-Acetyl-D-glucosamine | - | + |
| Turanose | - | - |
| Erythritol | - | + |
| Salicin | - | + |
| L-Glutamine | - | + |
| α-methyl-D-glucoside | - | NA |
| Glycyl-L- Proline | - | NA |
| Glycyl-L- Glutamine | - | - |
| Dextrin | + | NA |
| Arbutin | + | NA |
| D-Saccharic Acid | + | NA |
| a-D-Glucose | + | NA |
| L-Valine | + | NA |
| D-Cellobiose | + | NA |
| L-Rhamnose | + | NA |
| Succinic Acid | + | NA |
| D-Galactose | + | - |
| D-Mannose | + | NA |
| Palatinose | + | NA |
| Uridine-5'- Mono- phosphate | + | - |
| L-Fucose | + | NA |
| D-Glucose- 6-Phosphate | + | NA |
| Lactulose | + | + |
| D-Trehalose | + | NA |
| Succinic Acid Mono-Methyl Ester | + | NA |
| b-Cyclodextrin | + | NA |
| D-Sorbitol | + | NA |
| 3-Methyl-D- Glucose | + | NA |
| D-Arabitol | + | NA |
| D-Glucosaminic Acid | + | NA |
| Adonitol | + | + |
| D-Melezitose | + | NA |
| D-Mannitol | + | NA |

Antimicrobial susceptibility of the strain was tested using agar diffusion method in BHI agar medium. One hundred microliters of overnight grown culture of the strain was plated in BHI agar medium and antibiotic discs were placed. Zone of inhibition by antibiotics was measured after 24 hours of anaerobic incubation to determine the antibiotic sensitivity of the strain. The strain was found to be resistant to tetracycline and streptomycin. In contrast, the strain was susceptible to chloramphenicol, ampicillin, erythromycin and novobiocin.

## Genome sequencing information

### DNA isolation

Strain SG772 was sub-cultured on BHI agar and incubated at 37°C anaerobically for 48 hours. Further, it was cultured in 5 ml of BHI broth for 24 hours. Isolation of DNA was performed following E.Z.N.A DNA isolation kit (Omega, Biotek) protocol. Initially, 500μl of bacterial broth was pelleted by centrifuging at 10,000xg for 1 min. The pellet obtained was lysed in tris EDTA buffer using lysozyme for 1 hour and protein was digested using proteinase K overnight at 55°C. Subsequently, the mixture was pelleted by centrifuging at 8,000Xg for 1 min and supernatant was spun in spin column followed by two washes of wash buffer. Finally, DNA was eluted using 50 μl of nuclease free water and kept at −20°C until use.

### Genome sequencing, assembly and annotation

The genome was sequenced using llumina MiSeq (Illumina Inc, CA) usong 2x 250 paired end chemistry. Further, it was assembled using SPAdes 3.9.0 (13) and validated using Quast (14). Coding sequences were predicted using Glimmer 3.0 (15) and annotated with RAST 2.0 server (16) and the NCBI Prokaryotic Genomes Annotation Pipeline (PGAP) version 4.4. For core genome analysis, GenBank files were retrieved for strains SG772 along with its RAST neighbor *Blautia hansenii* DSM 20583 and were used for core genome analysis using OrthoMCL clustering algorithm (17) at 75% query coverage and sequence similarity (18). For functional annotation, the amino acid sequences were searched against the COG database to predict the abundance of COG gene families and heat map was constructed using Pearson correlation method and hierarchical clustering.

### Genomic attributes

Assembly of the *B. brookingsii* SG772 genome produced 55 contigs with the genome size of 3.46 Mbp (N50 = 179,176) and 43.97% GC content (Table 2). The largest contig was of 401,827 bp while the smallest was of 624 bps. The genome contains 57 tRNA genes, 3343 coding sequences, a phage element and a CRISPR element (Table 2).

**Table 2:** General attributes of *Blautia brookingsii* SG772 with *Blautia hansenii* DSM20583

| Characteristics | *Blautia brookingsii* SG772 | *Blautia hansenii* DSM20583 |
| --- | --- | --- |
| Genome size (bp) | 3,459,071 | 3,065,946 |
| Number of contigs | 55 | 1 |
| Coding sequences | 3343 | 3073 |
| %GC content | 43.97 | 39.04 |
| tRNA | 57 | 65 |
| rRNAs (5S, 16S, 23S) | 1, 8, 7 | 5, 5, 5 |
| ncRNAs | 4 | 4 |
| Pseudogenes | 152 | 68 |
| CRISPR elements | 1 | 2 |
| Phage | 1 | 0 |

## Insights from genome sequence

### Insight into the genetic repertoire

In order to compare the newly sequenced genome of strain SG772 to its closest RAST neighbor
*B. hansenii* DSM 20583 (8), we used BLAST Ring Image Generator (BRIG) (19) as a tool to show genome wide sequence similarity using *B. hansenii* DSM 20583 as a reference genome (Figure 4). Based on average nucleotide identity, these two genomes were 81.69% identical (19). Furthermore, this comparison revealed several differences in genome size, GC content and RNA copies (Table 2). For functional annotation, the amino acid sequences were searched against the COG database to predict the abundance of COG gene families and were compared against *B. hansenii* DSM 20583 (21). The gene families for energy production and conversion (C), amino acid transport and metabolism (E), carbohydrate transport and metabolism (G), transcription, ribosomal structure and biogenesis (J), transcription (K), replication, recombination and repair (L), cell wall/membrane/envelope biogenesis (M), inorganic ion transport and metabolism (P), defense mechanism (V) and unknown functions (S) were differentially enriched among these two strains (Figure 3C). Core genome analysis revealed a total of 411 core genes coding for basic metabolic functions. A total of 4,880 COG functions were annotated out of which only 70% were assigned to known categories. Additionally, resistance to cadmium, tetracycline, vancomycin, beta lactamase and fluoroquinolones along with dormancy and sporulation genes were relatively abundant in strain SG772. Furthermore, the genome was devoid of genes for motility and chemotaxis suggesting its non-motile phenotype.

**Figure 4:**
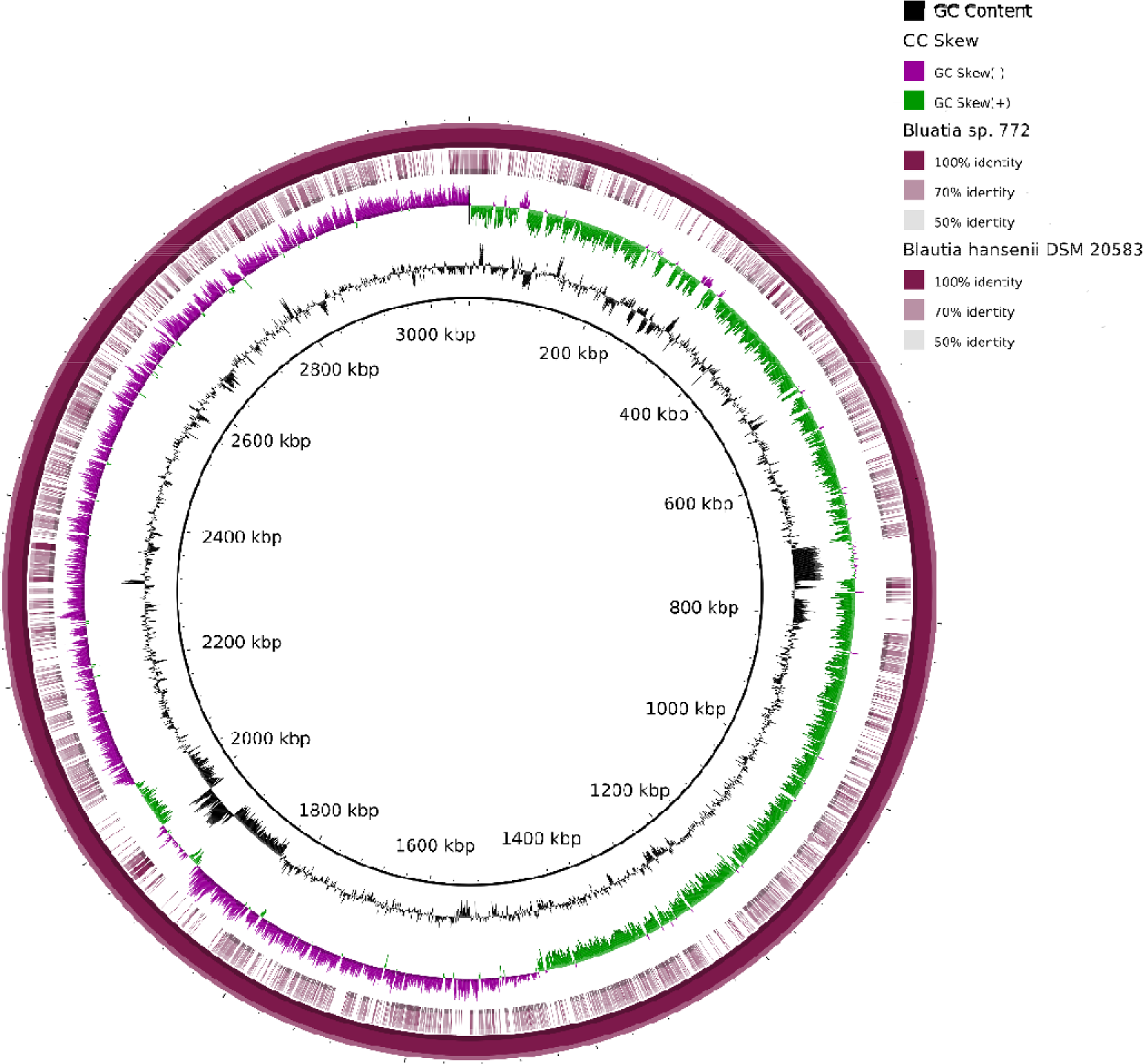
Comparative genome map of *Blautia sp.* SG772 using *B. hansenii* DSM 20583 as a reference. From the inside out, circle 1 represents the mean centered G+C content; circle 2 shows GC skew calculated as (G−C) / (G+C); circle 3 represents the genome sequence similarity of *Blautia sp.* SG772 based on color gradient against the reference genome of *B.xhansenii* DSM 20583; while circle 4 represents the genome of *B. hansenii* DSM 20583.

## Conclusions

This study presents the genome sequence for the strain SG772, with marked physiological and genomic differences from its neighbors *i.e*., *B. stercoris* GAM6-1 and *B. hansenii* DSM20583. Based on the differences, we propose a novel species of genus *Blautia* named as *Blautia brookingsii* SG772.

## Taxonomic and nomenclatural proposals

*Blautia brookingsii* (*brookingsii* referring to the isolation site of the type strain from Brookings, SD, USA). The cells are Gram stain positive, non-motile, coccobacillus, and 1.8–2.5 μm in length and 0.5–0.8 μm in diameter. Colonies were whitish, circular, smooth, and convex after 48 h of incubation in BHI agar. Optimal growth was observed at 37°C and pH of 6.8. It assimilates D-Mannitol, D-Melezitose, Adonitol, D-Glucosaminic acid, D-Arabitol, D-Sorbitol, D-Trehalose etc. Furthermore, the strain was resistant to tetracycline and streptomycin antibiotics whereas, it was susceptible to chloramphenicol, erythromycin, novobiocin and ampicillin. Genome sequence revealed the genome size of the strain SG772 to be 3.46Mbp with G+C content of 43.97 %. The type strain SG772 isolated from hlthy human fecal sample collected at Brookings, SD, USA and has been deposited in the Microbial Culture Collection at the National Centre for Cell Science, India.

## Declarations

Authors declare no competing interests.

## Acknowledgements

This work was supported in part by the USDA National Institute of Food and Agriculture, Hatch projects SD00H532-14 and SD00R540-15 to JS, and a grant from the South Dakota Governor’s Office of Economic Development awarded to JS, EN and JH.

